# Evaluation of InSeq to Identify Genes Essential for *Pseudomonas aeruginosa* PGPR2 Corn Root Colonization

**DOI:** 10.1101/377168

**Authors:** Ramamoorthy Sivakumar, Jothi Ranjani, Udayakumar S. Vishnu, Sathyanarayanan Jayashree, Gabriel L. Lozano, Jessica Miles, Nichole A. Broderick, Changhui Guan, Paramasamy Gunasekaran, Jo Handelsman, Jeyaprakash Rajendhran

## Abstract

The reciprocal interaction between rhizosphere bacteria and their plant hosts results in a complex battery of genetic and physiological responses. In this study, we used insertion sequencing (INSeq) to reveal the genetic determinants responsible for the fitness of *Pseudomonas aeruginosa* PGPR2 during root colonization. We generated a random transposon mutant library of *Pseudomonas aeruginosa* PGPR2 comprising 39,500 unique insertions and identified genes required for growth in culture and on corn roots. A total of 108 genes were identified as contributing to the fitness of strain PGPR2 on roots. The importance in root colonization of four genes identified in the TnSeq screen was verified by constructing deletion mutants in the genes and testing them for the ability to colonize corn roots singly or in competition with the wild type. All four mutants were affected in corn root colonization, displaying 5-to 100-fold reductions in populations in single inoculations, and all were outcompeted by the wild type by almost 100-fold after seven days on corn roots in mixed inoculations of the wild type and mutant. The genes identified in the screen had homology to genes involved in amino acid catabolism, stress adaptation, detoxification, signal transduction, and transport. INSeq technology proved a successful tool to identify fitness factors in P. *aeruginosa* PGPR2 for root colonization.

## Introduction

The rhizosphere is a dynamic, nutrient-rich environment on and around roots that supports intense activity among the resident soil microbiota and with the plant host. Root colonization is a complex process influenced by many factors. The primary bacterial traits important in root colonization are the ability of the bacterium to compete for niche space and respond to nutrients in the environment. This process involves sensing, response regulation, and chemotaxis toward the nutrient source (Broek et al., 1995). In some soil microbes, such as *Pseudomonas* sp., initial adhesion of bacteria to the root surface has been shown to trigger the expression of cell density-regulated genes that shape community behavior such that there is a direct correlation between bacterial population density and seed colonization (Espinosa-Urgel et al., 2004). This coordinated gene expression enables the bacteria to establish as microcolonies (multicellular aggregates) and eventually leads to the formation of biofilm-like structures on the root surface. Previous work has identified other bacterial traits that contribute to root colonization by bacteria, such as synthesis of amino acids, uracil, and vitamin B1, site-specific recombinase Sss, NADH dehyrogenase, and a Type Three Secretion System (TTSS) (Lugtenberg and Kamilova, 2009). Moreover, soil bacteria that positively influence the growth of plants, termed plant-growth-promoting rhizobacteria (PGPR), trigger a cascade of molecular signals that play a vital role in establishing a mutualistic relationship. Insight into these signaling processes is essential to understanding this complex relationship and improving such beneficial interactions for the betterment of agricultural plant production (Morrissey et al., 2004).

Fluorescent pseudomonads are well-known for their beneficial associations with plants derived from their aggressive colonization of roots and production of antimicrobial compounds active against pathogens (de Weger et al., 1986; Chin-A-Woeng et al., 2000; Lugtenberg et al., 2001). However, roles of plant root-derived compounds and the suite of genes involved in these beneficial associations have not been fully elucidated (Costa et al., 2007; Hartmann et al., 2009). In this study, we report the molecular determinants of *Pseudomonas aeruginosa* PGPR2 that are essential for root colonization of corn. Although *P*. *aeruginosa* has often been reported as an opportunistic pathogen of humans, it is also found in association with plants and some strains promote plant growth (Anjaiah et al., 2003; Walker et al., 2004).

*P. aeruginosa* PGPR2 is an efficient root colonizer and promotes plant growth. Previously, we reported the antagonistic properties of this bacterium against *Macrophomina phaseolina*, a fungal pathogen of plants (Illakkiam et al., 2013). Sequence analysis of the P. *aeruginosa* PGPR2 genome identified genes that might contribute to plant-growth promotion and disease suppression, including genes responsible for ACC deaminase, activation of auxin signaling, siderophore production, antifungal compound synthesis (i.e. HCN, phenazines), and phosphate solubilization (Illakkiam et al., 2014). However, whether these genes directly contribute to the proposed functions is not known.

Next-generation sequencing technology has opened a new area in functional genomics research. High throughput transposon insertion sequencing, such as HITS (Gawronski et al., 2009), TraDIS (Langridge et al., 2009), INSeq (Goodman et al., 2009) and Tn-seq (van Opijnen et al., 2009), has emerged as a powerful functional genomics tool that establishes causal relationships between genes and bacterial behavior. This strategy combines transposon mutagenesis and high-throughput sequencing, which allows simultaneous assessment of the fitness of thousands of discrete mutants for a particular function. Thus, INSeq enables identification of genetic elements required for the fitness of an organism *in vitro* or *in vivo*. Recently, Cole et al. (2017) reported *Pseudomonas simiae* genes required for *Arabidopsis thaliana* root colonization using randomly barcoded transposon mutagenesis sequencing (RB-TnSeq). In the study reported here, we employed INSeq analysis to unravel the genetic elements responsible for mutualistic interactions using corn and *P. aeruginosa* PGPR2 as a model system. We identified 108 genes that were essential for fitness of strain PGPR2 during corn root colonization.

## Material and Methods

### Bacterial strains and growth conditions

The bacterial strains and plasmids used in this study are listed in Table 1. *Pseudomonas aeruginosa* PGPR2 and *Escherichia coli* were grown at 30°C and 37°C, respectively, and routinely sub-cultured in Luria Bertani (LB) medium. When required, the medium was solidified using 2% agar. Antibiotics were added as needed at the following concentration (unless otherwise specified): ampicillin, 100 μg ml^−1^; gentamicin, 40 μg ml^−1^; irgasan, 25 μg ml^−1^.

### Construction of pSAM-BT20 vector

The transposon delivery vector, pSAM-BT20, was derived from pSAM-BT (Goodman et al. 2009). The erythromycin resistance gene was removed with *XhoI* and *XbaI* (New England Biolabs, Ipswich, MA) and replaced with a gentamicin resistance gene that was PCR amplified with primers GenF-GenR (Table S1). The transposase gene was removed with *BamHI* and *NotI*(New England BioLabs) and replaced with the transposase gene from pBT20, which was PCR amplified using primers Trans F-Trans R (Goodman et al., 2009).

### Construction of transposon insertion mutant library

The transposon library was constructed by conjugation of *P*. *aeruginosa* PGPR2 with *E. coli* λ *pir* harboring pSAM-BT followed by selection of exconjugants for gentamicin resistance. Briefly, the donor and recipient strains were grown separately overnight. Cultures were mixed and pelleted, then washed with fresh LB medium, suspended in 100 μl of LB medium and spotted on an LB agar plate, and incubated at 30°C for 3 hours. The exconjugants containing insertions were selected by plating on LB medium supplemented with gentamicin (40 μg ml^−1^) and irgasan (25 μg ml^−1^) for counterselection against the donor *E. coli* strain. The plates were incubated at 30°C for 24 hours. Successful integration of the transposon into the genome was verified by performing PCR for the gentamicin resistance gene using primers GenF and GenR. The colonies were pooled using sterile phosphate-buffered saline containing 15% glycerol. One ml aliquots of the mutant library suspension were placed in vials and stored at −80°C until further use. One vial was retrieved from the stock and enumerated by standard dilution plating.

Southern blot hybridization was performed to confirm random integration and single-copy insertion of the transposon into the genome of PGPR2. Genomic DNA was isolated from 13 random mutants using the Qiagen blood and tissue mini kit according to the manufacturer’s instructions. Genomic DNA (1 μg) was digested with *Hin*dIII and separated by electrophoresis on a 0.8% Tris-boric acid-EDTA agarose gel and transferred to a positively charged nylon membrane. The gentamicin-resistance cassette was used as a probe and labeled using the Digoxigenin-dUTP (DIG) DNA labeling kit (Roche, Switzerland) according to the manufacturer’s instructions. Bound DNA was hybridized with a DIG-labeled gentamicin-resistance gene generated by random primer amplification and visualized by autoradiography.

We generated a library of 39,500 insertion mutants in which integration of transposons into the genome was confirmed by PCR amplification from the genomic DNA of 20 random mutants, which showed the presence of the gentamicin resistance cassette, but not the transposase gene (Fig. S1). Southern blotting showed that each PGPR2 mutant had only one unique transposon insertion in the genome, as only one fragment from each mutant hybridized to the probe. Also, the single integration was random, as indicated by the range of size of the fragments carrying the transposon (Fig. S2). On average, transposon integration was found every 169 bp in the genome. Fig. S3 depicts the random insertions throughout the genome.

### Plant system for InSeq selection

Surface-disinfected corn seeds were germinated on moist filter paper for three days and transferred to the gnotobiotic hydroponic plant nutrient medium (Hoagland and Arnon, 1950). An aliquot of the mutant library was thawed, pelleted, and washed three times with sterile 10 mM MgSO_4_. The suspension was diluted to ~4 x 10^6^ CFU ml^−1^ and inoculated onto germinated seedlings in triplicate (n=3). A portion of the suspension was used for genomic DNA isolation (input pool). The plants were maintained in the greenhouse with 16 hours light and 8 hours of dark. Seven days after inoculation, the roots were aseptically excised, washed gently to eliminate weakly adhered bacterial cells and transferred to 50-ml tubes containing 0.85% saline and ten glass beads (3 mm in diameter). The tubes were vortexed briefly to detach bacteria from the root surface, the root was discarded and the suspension was used for genomic DNA isolation (output pool). The workflow is shown in Fig. 1.

**Fig. 1.**
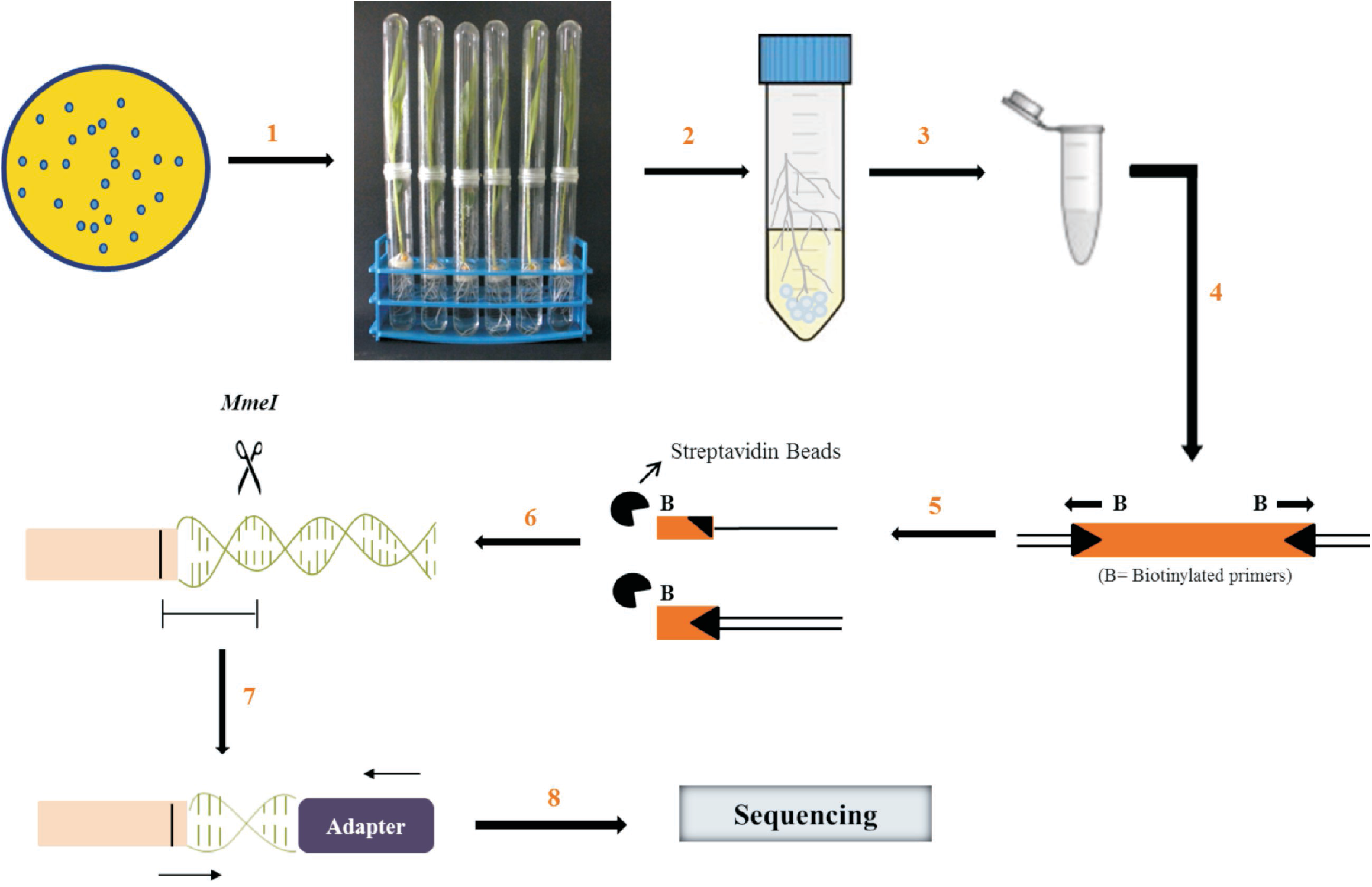
Workflow of insertion sequencing. A transposon insertion library in *P. aeruginosa* PGPR2 with ~40,000 mutants was generated and a suspension of 4 × 10^6^ CFU ml^−1^ was applied to germinated corn seedlings (step 1). Seven days post inoculation, the roots were excised aseptically and briefly vortexed in 20 ml saline containing glass beads to detach the bacteria adhered to root surface (step 2). Genomic DNA was extracted and purified from both input and output populations (step 3). Linear PCR was performed using biotinylated primers to amplify transposon integration site (step 4). The amplified product was captured using streptavidin beads and the second DNA strand was synthesized (step 5). A DNA fragment was digested with *MmeI* enzyme and ligated with Illumina barcodes (step 6). The product was amplified for a restricted number of PCR cycles and ligated with Ion Torrent adapters on either side (step 7). The sequencing was performed using the Ion Torrent platform (step 8) (adapted from Goodman et al., 2011).

### InSeq library preparation and sequencing

Total DNA was isolated from the input and output populations using QIAGEN DNeasy blood and tissue kit according to the manufacturer’s instructions. The extracted DNA was used as template to amplify transposon-insertion junctions with appropriate barcodes as previously described (Goodman et al., 2011). The amplicons obtained were ligated with Ion Torrent adapters, and samples were pooled in equimolar concentration and sequenced using Ion Torrent PGM on a 318 chip.

### Data analysis

Adapters were trimmed from the raw reads using Cutadapt and the reads were split based on barcodes using FastX barcode splitter. The INSeq data were analyzed using the online software ESSENTIALS (http://bamics2.cmbi.ru.nl/websoftware/essentials) (Zomer et al., 2012). The processed reads were mapped to the *P*. *aeruginosa* PGPR2 genome (Accession number ASQO01000001-ASQO01000198).

### Construction of *P. aeruginosa* PGPR2 deletion mutants

*P. aeruginosa* PGPR2 deletion mutants were generated in the *trpD, hom, oprF* and *cbrA* genes. Briefly, PCR primers were designed to amplify a partial fragment of each gene in the 5’ and 3’ regions and contained *HindIII* restriction sites (Table S1). The gentamicin-resistance cassette was amplified from the pUCP24 vector (Table 2) with primers containing *Hind*III restriction sites. Appropriate DNA fragments were then restricted and ligated with a gentamicin cassette resulting in deletions of each gene. This *in vitro* product was then cloned into the *Xho*I site of suicide vector pIVETP (Table 1), which was then mobilized into P. *aeruginosa* PGPR2 from E. *coli* S-17 *λ-pir*. The exconjugants resulting from double homologous recombination were selected on LB agar containing gentamicin and irgasan. Deletion alleles within each mutant were confirmed by PCR.

### Root colonization assay

To verify the reduction in colonizing ability of isogenic mutants constructed in *P. aeruginosa* PGPR2, root colonization assays were performed. Briefly, corn seeds were surface sterilized as above and germinated seedlings were transferred to gnotobiotic hydroponic medium. Corn plantlets grown in hydroponic system was inoculated with ~4 x 10^6^ CFU of wild type and mutant strains. At 7 days post inoculation, the roots were aseptically excised and transferred to a 50 ml tube containing 20 ml 0.85% saline. The tube was vortexed for 1 min to detach adhered bacteria, and the suspension was enumerated by standard dilution plating. The experiment was repeated three times with five replicates. Differences were considered statistically significant when p<0.001.

### Competitive root colonization

To test the competitive root-colonizing ability of mutant strains, 1:1 mixtures of the wild-type and each mutant strain were applied to plantlets grown in hydroponic medium as described above. After seven days, bacterial cells were isolated from the root surface and plated on LB medium supplemented with and without gentamicin to differentiate the mutant from the wild-type strain. A strain’s competitiveness was expressed as the percentage of total colony forming units recovered from corn roots represented by the strain. Each experiment was repeated three times with five replicates per treatment.

## Results

### INSeq genetic analysis of root colonization by *P. aeruginosa* PGPR2

We generated and validated a library of approximately 39,500 insertion mutants in *P. aeruginosa* PGPR2, which contains 6,803 genes, providing approximately 6-fold coverage or an average of six insertions per gene. To identify *P. aeruginosa* PGPR2 genes essential for the root colonization in corn, we inoculated each corn seedling with 4 x 10^6^ CFU of the insertion mutant library. The bacterial population increased 100-fold in seven days, after which the input and output populations were analyzed.

### Colonization mutants of *P. aeruginosa* PGPR2

A total of 993 genes were necessary for the growth of PGPR2 in LB medium, including genes required for normal growth, such as those encoding tRNA and rRNA (Table S2). The genes essential for the fitness of PGPR2 during root colonization were identified by comparing the number of reads for each insertion site in the output pool to the number of reads in the input pool. Mutants in 108 genes (2% of the predicted genes in PGPR2) were underrepresented in the output pool by at least two-fold (log_2_ < −1 with an adjusted p-value < 0.05) and therefore considered essential for fitness of the organism during root colonization (Fig. 2A and 2B) (Table S3). Most of these genes encoded functions associated with amino acid transport and metabolism, energy production and conservation, and coenzyme transport and metabolism. Other genes implicated in root colonization were associated with cell motility, nucleotide transport and metabolism, cell wall/membrane/envelope biogenesis, and signal transduction (Fig. S5).

**Fig. 2.**
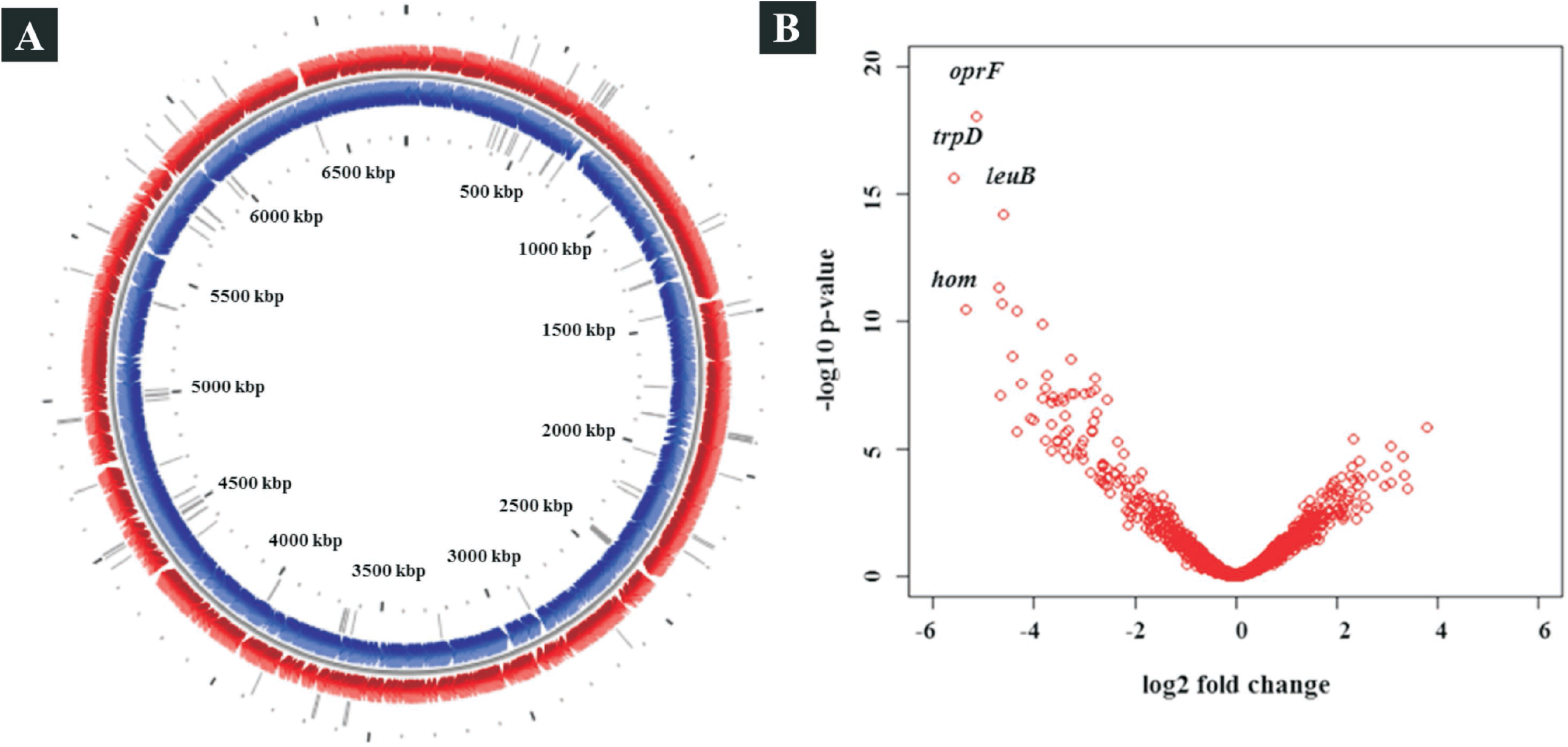
Mutants affected for colonization of corn roots. (A) Genome map showing transposon insertion sites of conditionally essential genes in *P. aeruginosa* PGPR2 genome. The outer circle represents forward strand (red), the inner circle represent reverse strand (blue), and the grey bars depict the transposon insertion sites. Circular plot was generated using the CG view tool (Stothard and Wishart, 2005). (B) Volcano plot representing the genes responsible for fitness of PGPR2 in the corn rhizosphere. The fitness genes with highest significance are highlighted with name designations.

### Validation of colonization mutant phenotype

To validate the INSeq experimental results, we constructed four isogenic mutants and tested for their ability to colonize corn roots both individually and in competition with the parent strain. We selected four genes, which showed the highest negative fold-changes (*trpD*, −48; *hom*, −41; *oprF*, −35; and *cbrA*, −11). All four knock-out mutants showed poor root-colonizing ability when tested either individually or when co-inoculated with the wild-type strain (Fig. 3), thereby validating the INSeq results.

**Fig. 3.**
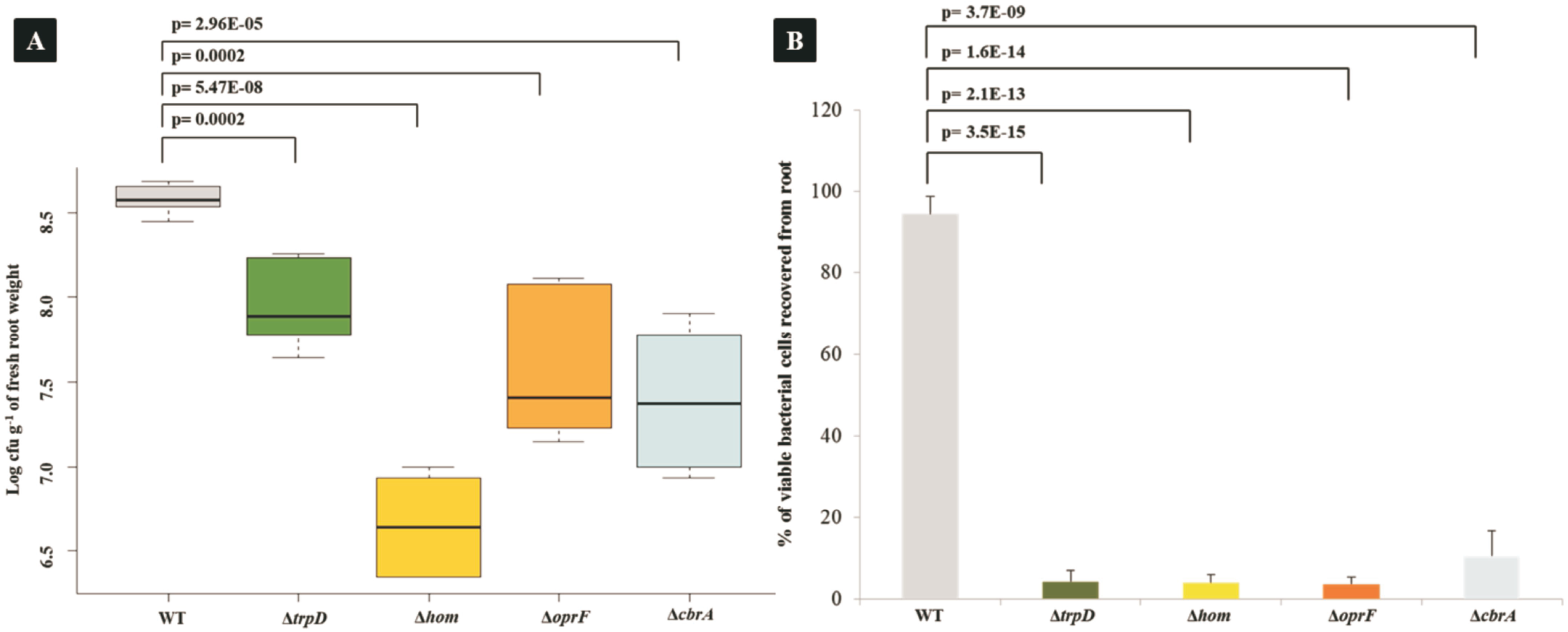
Validation of colonization mutants identified by INSeq screen. (A) Corn root colonization by wild type *P. aeruginosa* PGPR2 and its knockout deletion mutants. ~4 x 10^6^ CFU ml^−1^ of wild type *P. aeruginosa* PGPR2 and each of the mutant strains (ΔtrpD::Gm^R^, Δhom::Gm^R^, ΔoprF::Gm^R^ and *ΔcbrA* ::Gm^R^) were inoculated individually and bacterial populations were determined seven days post inoculation. The colonization experiment was performed thrice independently with five replicates within each experiment. (B) Competitive corn root colonization by wild-type *P. aeruginosa* PGPR2 and its deletion mutants. 1:1 mixtures of wild type and one of the mutants were inoculated; the competitive colonizing ability was expressed as percentage of viable bacterial cells recovered from root.

## Discussion

An InSeq mutant screen was performed in P. *aeruginosa* PGPR2, a plant-growth promoting bacterium and an efficient root colonizer, to identify genes involved in root colonization. We found that a total of 993 genes are essential for the growth in LB medium. Functional categorization revealed many of these essential genes are involved in translation (9%), transcription (8.3%), replication (7.8%), cell wall biogenesis (8.5%) and genes with unknown function (20.5%) (Fig. S4). Earlier, Jacob et al., (2003) reported that 678 genes were essential for growth of *P. aeruginosa* PAO1 in LB medium. In another study, Liberati et al., (2006) identified 1454 genes that were essential for the growth of *P. aeruginosa* PA14, using an ordered, non-redundant transposon mutant library.

We found that 108 genes of PGPR2 are conditionally essential for the corn root colonization. Functions of these genes, their possible roles in root colonization, and results from other studies that identified genes required for root colonization are discussed below.

### (i) Motility and adhesion

The bacterial fitness and colonization of plant surfaces during plant-microbe interaction is directly associated with motility owing to its role in nutrient access, avoidance of toxic materials, niche competition with other microorganisms, host preference, and maneuvering efficiently in the rhizosphere (Nogales et al., 2015). Transposon insertion in two genes involved in motility, *flhF* and *flgD*, led to a significant (3.7-to 8.1-fold) reduction in fitness (Table 2).

### (ii) Energy production

NADH dehydrogenase or NADH/ubiquinone oxidoreductase encoded by the *nuo* operon is a key enzyme in energy production responsible for electron transfer to one of the two terminal oxidase complexes. Of the fourteen genes comprising this operon, our screen found that transposon insertion in seven genes *(nuoH, nuoB, nuoK, nuoE, nuoN, nuoM* and *nuoG)* led to a 10.4-to 4.4-fold reduction in fitness (Table 2). A previous study showed that mutations in NADH dehydrogenase resulted in 100-fold impairment of colonization efficiency of *P. fluorescens* WCS365 (Margarita et al., 2002). Three genes involved in cytochrome biosynthesis, encoding cytochrome C assembly protein, cytochrome B and cytochrome oxidase subunit I were significantly underrepresented in the output pool from corn roots. The mutants were impaired in colonization and displayed a fitness reduced from 4.5-to 3.0-fold (Table 2), in agreement with a previous study that reported overexpression of cytochrome C oxidase genes in the rhizosphere during colonization by *P. putida* KT2440 (Matilla et al., 2007).

### (iii) Carbon metabolism

Transposon insertions in two signature genes of the glyoxylate cycle, isocitrate lyase and malate synthase, were underrepresented in the population with the fold-change of −7.3 and −6.8 (Table 2). The glyoxylate cycle is a key carbon anaplerotic pathway that bypasses the decarboxylative steps of tricarboxylic acid (TCA) cycle to produce malate and succinate by utilizing two molecules of acetyl-CoA when generation of pyruvate from the glycolysis cycle is prevented (Dunn et al., 2009; Fleck et al., 2011). In a previous study, Ramos-Gonzalez and coworkers (2005) showed that isocitrate lyase synthesis was induced in *P. putida* KT2440 during corn root colonization. The role of these genes in colonization might be due to acetate providing a major carbon component in the plant exudate (Lugtenberg and Bloemberg, 2004), which is acted upon by isocitrate lyase, converted to acetyl-coenzyme A (CoA), and subsequently utilized for energy production.

### (iv) Nitrogen assimilation

Nitrogen assimilation is a delicate balancing act mediated by *ntr* and *gln* systems in Proteobacteria. Transposon insertions in *glnD* significantly reduced the fitness of PGPR2 during root colonization with the negative selection of −5.8 fold (Table 2). GlnD is a bifunctional uridylyltransferase/uridylyl removing enzyme that regulates PII proteins by uridylylation or deuridylylation and is thought to be the principal regulator of intracellular nitrogen status under nitrogen-deprived conditions (Zhang et al., 2005). Mutations that affect regulatory *gln* genes result in defective nitrogen acquisition during both free-living and symbiotic states and *glnD* mutants of *Sinorhizobium meliloti* do not form normal symbiotic interactions with alfalfa (Yurgel and Kahn, 2008).

### (v) Amino acid metabolism

Our screen confirmed a previous report of the importance of amino-acid synthesis for root colonization (Simons et al., 1997), (Fig. 4). We found transposon insertions in four genes involved in tryptophan biosynthesis that exhibited 11-to 48-fold reductions in fitness (Table 2). Bioavailability of metabolites such as amino acids in root exudate may support colonization of plant roots by *P. putida* (Matilla et al., 2007). Tryptophan is the precursor for indole-3-acetic acid (IAA), a major plant hormone that can trigger growth and several other responses in the plant (Spaepen et al., 2007 and Roca et al., 2012). Tryptophan is a costly amino acid used by bacteria as a building block for the production of siderophores, signaling molecules, and disease-suppressive compounds (Matthijs et al., 2004; Farrow et al., 2007 and Balibar et al., 2006), which are essential for root colonization.

**Fig. 4.**
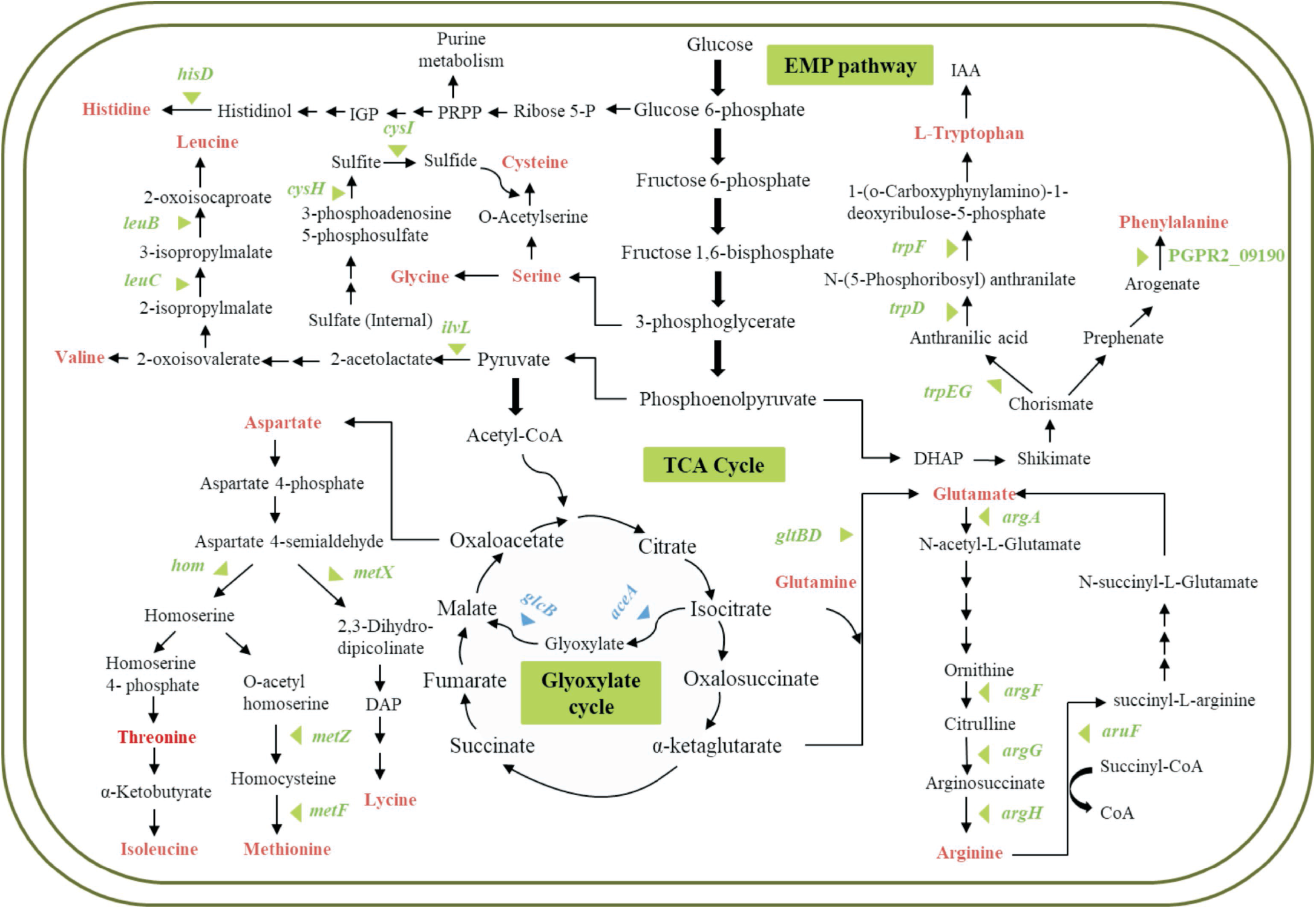
Essential genes involved in metabolic pathways (carbon and amino acid metabolism) responsible for fitness of *P. aeruginosa* PGPR2 in the corn root. Genes significantly (q<0.05) influenced fitness during root colonization are marked with triangles; blue triangles indicate carbon metabolism and green triangles indicate amino acid metabolism.

The transposon insertion in a gene involved in the metabolism of aspartate resulted in a 41-fold reduction. This gene codes for homoserine dehydrogenase, which is upregulated in *Rhizobium leguminosarum* during interaction with pea (Ramachandran et al., 2011). Similarly, transposon insertions in genes involved in arginine and methionine biosynthesis pathways significantly reduced fitness of PGPR2 from 13.3-to 4.4-fold during root colonization. Similarly, transposon insertions in the genes involved in the metabolism of leucine, histidine, isoleucine, glutamate and cysteine reduced fitness from 24.7-to 3.5-fold. Similarly, *P. fluorescens* WCS365 mutants auxotrophic for the amino acids arginine, leucine, histidine, and isoleucine have greatly diminished root-colonizing ability (Simons et al., 1997), while genes involved in isoleucine biosynthesis are induced in the sugar beet rhizosphere during colonization of *P. fluorescens* SBW25 (Gal et al., 2003).

### (vi) Nucleotide metabolism

We identified three genes involved in purine biosynthesis (*purF* coding for amidophosphoribosyl transferase, *purL* coding for phosphoribosyl formylglycinamidine synthase, and *purM* coding for phosphoribosyl aminoimidazole synthetase) as being significantly underrepresented after root colonization. Mutants in these genes were significantly hampered in root colonization ability, reduced 8.4-to 10.5-fold (Table 2). Similarly, previous work showed that a *purM* mutant of *P. chlororaphis* O6 failed to neither colonize tobacco roots nor induce systemic resistance (Han et al., 2006).

### (vii) Fatty acids and vitamin metabolism

Transposon insertions in two genes involved in fatty acid metabolism coding for acyl-CoA dehydrogenase and enoyl-CoA hydratase reduced fitness on roots 3.7-to 13.5-fold (Table 2). This result reinforces an earlier study (Cheng et al., 2009) that reported the acyl-CoA dehydrogenase gene of *P. putida* UW4 is over-expressed during interaction with *Brassica napus*. Similarly, *Bacillus amyloliquefaciens* FZB42 over-expressed an enoyl-CoA hydratase in response to corn root exudates (Fan et al., 2012).

Vitamins are essential micronutrients synthesized by most plants and bacteria (Survase et al., 2006) and function as cofactors for many cellular activities and confer protection against free radicals (Asensi-Fabado and Munné-Bosch, 2010 and Smith et al., 2007). The role of vitamins is vital, especially in plant-microbe interactions, as they promote the growth of PGPRs around the root system (Palacios et al., 2014). Transposon insertions in *cbiA, thiG, bio,B* and *pdxA*, which are involved in the synthesis of water soluble vitamins, reduced fitness from 4.8-to 5.2-fold (Table 2). *thiG* codes for thiazole synthase, which couples with pyrimidine to synthesize thiamine (Begley et al., 1999). Thiamine is an essential vitamin acting as a cofactor for the enzyme indolepyruvate decarboxylase, which plays a role in the synthesis of IAA in some PGPRs. *bioB* codes for biotin synthase, which is involved in the synthesis of vitamin B7. Streit and colleagues (1996) reported that biotin promotes root colonization of *Rhizobium meliloti* on alfalfa roots. The *pdxA* gene codes for 4-hydroxythreonine-4-phosphate dehydrogenase, an enzyme involved in the synthesis of vitamin B6 (pyridoxine), a vitamin vital for protection of bacterial cells from oxidative stress (Asensi-Fabado and Munné-Bosch, 2010).

### (viii) DNA recombination and repair mechanism

Genetic rearrangements and horizontal DNA transfer events are frequent in the rhizosphere (Matilla et al., 2007). Genes involved in these processes such as *ruvB* encoding a DNA helicase, *parA*, encoding a chromosome-partitioning gene, and an insertion sequence ISPpu14 transposase were found to be important for root colonization by PGPR2. Disruption of these genes resulted in decreased fitness with the fold-change of 4.5-to 19-fold (Table 2). This is in agreement with Matilla et al., 2007 who found that *P. putida* KT2440 helicase and transposase ISPpu14 are upregulated in the rhizosphere.

### (ix) Stress adaptation and detoxification

Three genes, *algU*, *gshA*, and *gshB*, involved in stress responses, were underrepresented in the output population with reductions between 3.4- and 16.8-fold (Table 2). The *algU* gene codes for the extracytoplasmic function (ECF) sigma factor, which is a transcriptional regulator triggered by environmental stimuli (Potvin et al., 2008). Extracellular polysaccharide production is mostly regulated by the sigma factor AlgU (also referred as AlgT) in the biocontrol agent *Pseudomonas fluorescens* CHA0 and is required for optimal survival of the strain under desiccation and osmotic stress conditions (Schnider-Keel et al., 2001). AlgU also activates genes responsible for the production of alginate in *P. syringae* and contributes to the *in planta* growth and survival of P. *syringae* pv. *glycinea* (Schenk et al., 2008). *gshA* codes for glutamate-cysteine ligase, which is involved in the biosynthesis of glutathione and is a key protectant against oxidative stress. Similarly, disruption of *gshB*, which codes for glutathione synthase, resulted in a competitive disadvantage of *Rhizobium tropici* during bean nodulation (Riccillo et al., 2000). Thus, the AlgU sigma factor and *gsh* genes might contribute to counteracting stress in the rhizosphere.

### (x) Sensors and regulator proteins

Two-component system (TCS) signaling pathways are the primary signaling mechanisms by which bacteria sense and respond to environmental cues. They are crucial for many adaptive mechanisms including motility, chemotaxis, and metabolism (Gao and Stock 2009). Transposon insertions in five genes involved in TCS reduced fitness from 4.6-to 10.8-fold (Table 2). One of these genes, *cbrA*, codes for a putative two-component histidine kinase is involved in the stress response of *Pseudomonas putida* KT2440 (Reva et al., 2006). Similarly, disruption of *cbrA* was reported to down-regulate genes responsible for motility, chemotaxis, metabolism, and cell wall function (Gibson et al., 2006 and 2007) and *P. aeruginosa* mutants lacking CbrA fail to utilize several carbon sources and amino acids (Nishijyo et al., 2001). We also identified a mutant contain a mutation in a gene encoding a hybrid sensor kinase RetS, which is a regulator of exopolysaccharide and type III secretion system, whose fitness was reduced 9.5-fold). The same gene is crucial in the temperature response of *P. fluorescens* CHAO (Humair et al., 2009).

Insertions in genes associated with the ArsC (PGPR2_08335) and MarR (PGPR2_12420) transcriptional regulators resulted in significantly reduced fitness by 8.9- and 5.8-fold (Table 2). Members of the multiple-antibiotic resistant regulator (MarR) are responsible for resistance to antibiotics, organic solvents, and oxidative stress agents (Alekshun and Levy, 1999).

### (xi) Transporters

Transposon insertion in four ABC transporters (PGPR2_29805, PGPR2_12725, PGPR2_25320 and PGPR2_29810) reduced colonization fitness with fold-changes from 6.2-to 3.0- (Table 2). These genes code for an ABC transporter permease, multidrug efflux pump, and VacJ ABC transporter. Matilla et al., (2007) reported the up-regulation of an ABC transporter permease during the interaction of *P. putida* KT2440 with corn, in which the plant root exudes a diverse array of secondary metabolites with antimicrobial activity (Dixon, 2001). Multidrug resistance (MDR) pumps have been shown to be important for phytopathogens to detoxify such antimicrobials (Maggiorani et al., 2006). *Sinorhizobium meliloti* with mutations in MDRs display reduced nodulation competitiveness due to sensitivity to antimicrobial compounds produced by the host plant (Eda et al., 2011).

Disruption of two genes coding for outer membrane proteins OprF and OprD resulted in reduced fitness with the fold-change of 35 and 3.3 (Table 2). OprF, the major outer membrane protein found in *Pseudomonas*, is responsible for transport of small molecules such as nutrients and waste products across the outer membrane and *P. fluorescens* strain OE28.3 mutants lacking OprF exhibited impaired adhesion to wheat roots (De Mot et al., 1993).

### (xii) Osmoregulation

The release of solutes by plant root during the uptake of water, discharge of root exudates, and the production of exopolysaccharides increases osmolality of the rhizosphere. Thus, the ability of rhizobacteria to adapt to elevated osmolality is essential for successful root colonization. This is achieved by tight regulation of the periplasmic glucan under variable osmolality (Miller and Wood, 1996). The cell envelope of Gram-negative bacteria among the Proteobacteria have osmoregulated periplasmic glucans (OPGs), which play a vital role during the growth under hypoosmolarity. Transposon insertions in genes involved in the biosynthesis of glucans were significantly underrepresented in our screen with fold-changes of −12.5 and −9.2 (Table 2). These genes code for a periplasmic glucan biosynthesis protein (MdoG) and glucosyltransferase (MdoH), which is required for virulence of plant pathogens, including *Erwinia chrysanthemi* and *Pseudomonas syringae* (Loubens et al., 1993; Page et al., 2001).

### (xiii) Protein folding and degradation

In order to survive and proliferate efficiently, bacteria require specific proteases that degrade misfolded proteins and regulatory proteins to cope with stress conditions such as those that occur during plant-microbe interactions. Proteolysis of such proteins is carried out by energy-dependent proteases, such as members of the Clp family (e.g., ClpAP and ClpXP). The Clp family proteases are essential for bacterial fitness as they are implicated in several stress responses. The primary function of these proteins is to govern the folding and refolding of stress-response proteins. A transposon insertion in *clpX*, which codes for ATPase subunit of the Clp protease (PGPR2_18895), significantly reduced fitness *in vivo* with the fold-change of 14 (Table 2). Mutation in this gene altered the biofilm-forming ability of *P. fluorescens* WCS365 (O’Toole and Kolter, 1998).

## Conclusion

We employed the transposon-insertion sequencing technique, INSeq, to identify fitness determinants of *P. aeruginosa* PGPR2 during root colonization of corn. We uncovered a total of 108 genes that contribute to root colonization by PGPR2 with many having functions previously shown to be important in root colonization, thus validating the INSeq approach. The key functions required for root colonization identified in this study are integrated and shown in Fig. 5. Further characterization of the fitness genes essential for root colonization may provide insights into the novel pathways employed by PGPRs in establishing mutualistic relationships with their hosts. This study sheds light on the power of genome-wide approaches like INSeq in understanding complex mechanisms underlying plant-microbe interactions. Deciphering these relationships paves the way to promote these beneficial interactions for improving crop productivity in agriculture.

**Fig. 5.**
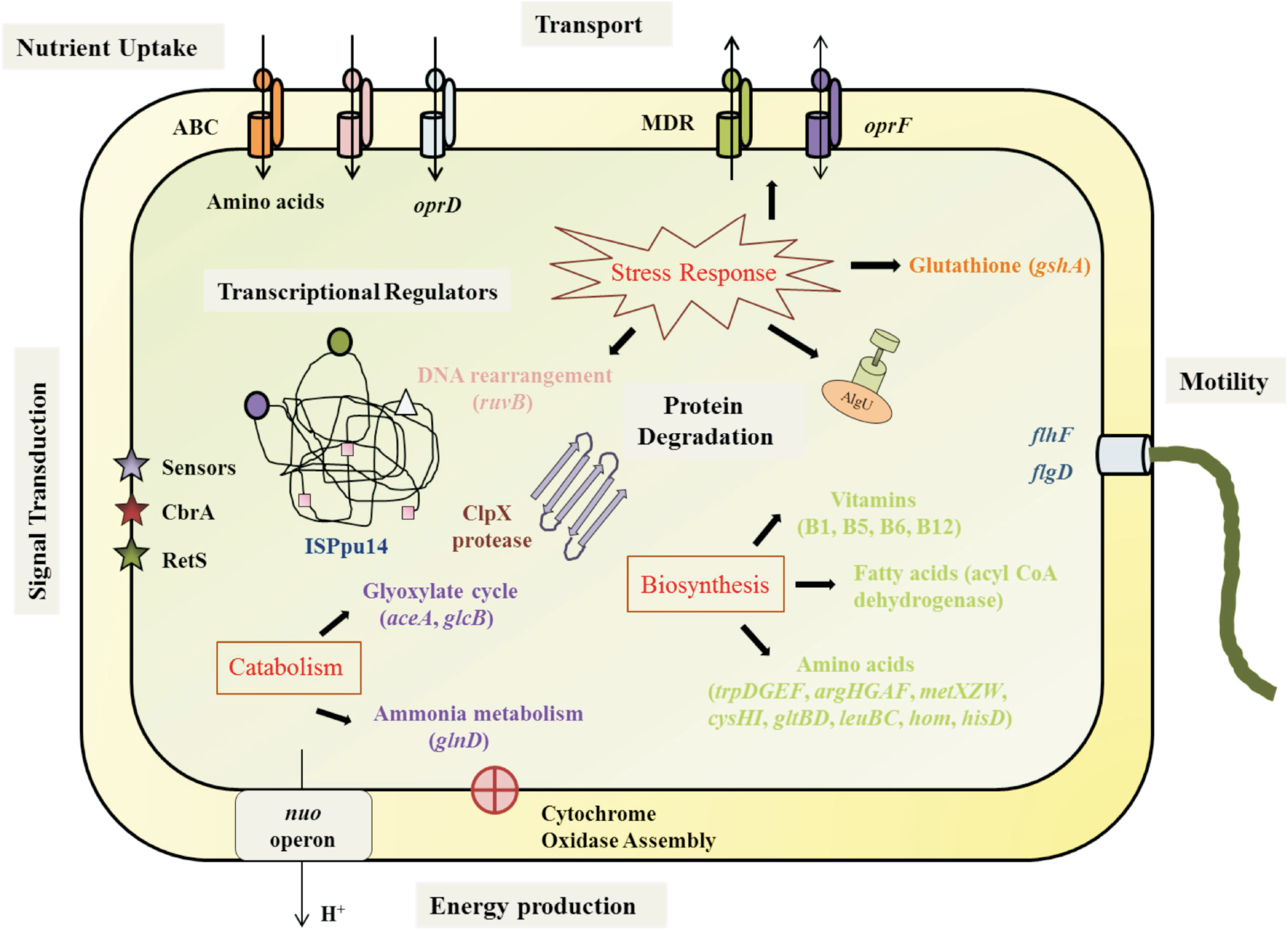
Integrated scheme showing the fitness genes identified in this study and their functions. See text for details

## Acknowledgements

RS acknowledges the Council of Scientific and Industrial research (CSIR) for providing Senior Research Fellowship (09/201/0416/2016-EMR-I). JR acknowledges the UGC-Raman Postdoctoral Fellowship (F.No. 5-38/2013 (IC)) from University Grants Commission, New Delhi. The UGC-CAS, NRCBS, DBT-IPLS, DST-PURSE Programs of the School of Biological Sciences, Madurai Kamaraj University is gratefully acknowledged. RS acknowledges Dr. Sridevi, Mr. Victorathisayam and Ms. Rajpriya, School of Biotechnology, Madurai Kamaraj University, Madurai for their help in performing Southern blotting. RS acknowledges Dr. Shankar Manoharan, Indian Institute of Technology Jodhpur, for his constructive suggestions. This work was also supported by NSF grant MCB-1243671.

## Authors Contributions

JR, PG, and JH conceived and designed the work. RS constructed the mutant library. RS and JoR constructed the library for sequencing. RS and USV performed sequencing. RS, USV, JoR, SJ, and JR analyzed and interpreted data. RS constructed the mutant strain for validation. GL, JM, NB, and CG constructed the plasmid used for transposition. RS, JR, PG, SJ, GL, JM, NB, CG, and JH wrote the manuscript.

---

**Fig. S1.**
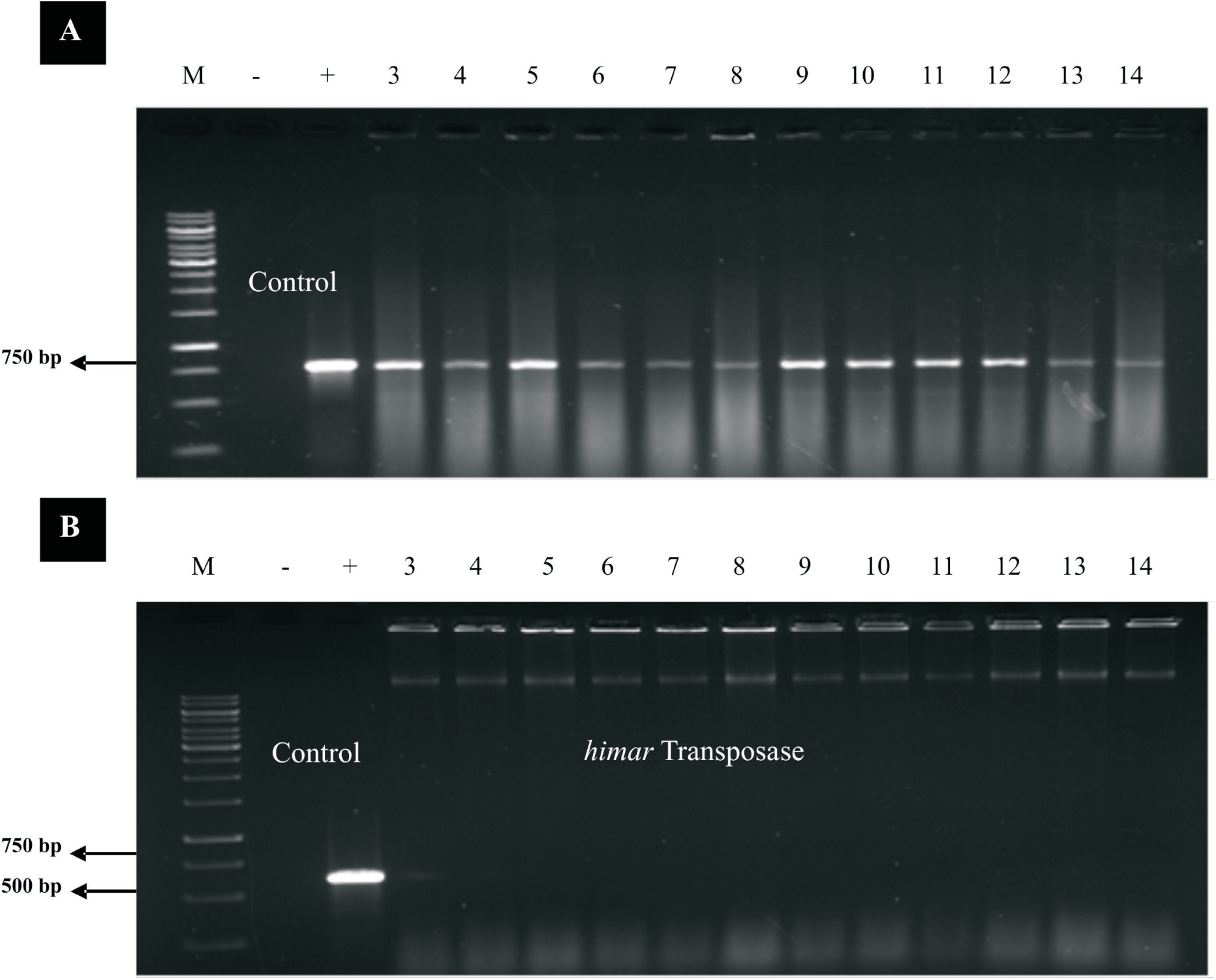
Confirmation of transposon integration in PGPR2 mutant library. (A) Integration of mariner transposon was confirmed with PCR to amplify the transposon. Lane M, 1-kb molecular weight DNA marker (Thermo Fisher Scientific, USA); “-”, negative control (gDNA from P. *aeruginosa* PGPR2); “+”, positive control (pSAM_BT plasmid DNA); lane 3-14, PCR amplification of Gm^R^ antibiotic cassette from gDNA of individual mutants of *P. aeruginosa* PGPR2. (B) PCR for the same samples using *himar* transposase gene primers should be lost in mutants upon proper integration of transposon.

**Fig. S2.**
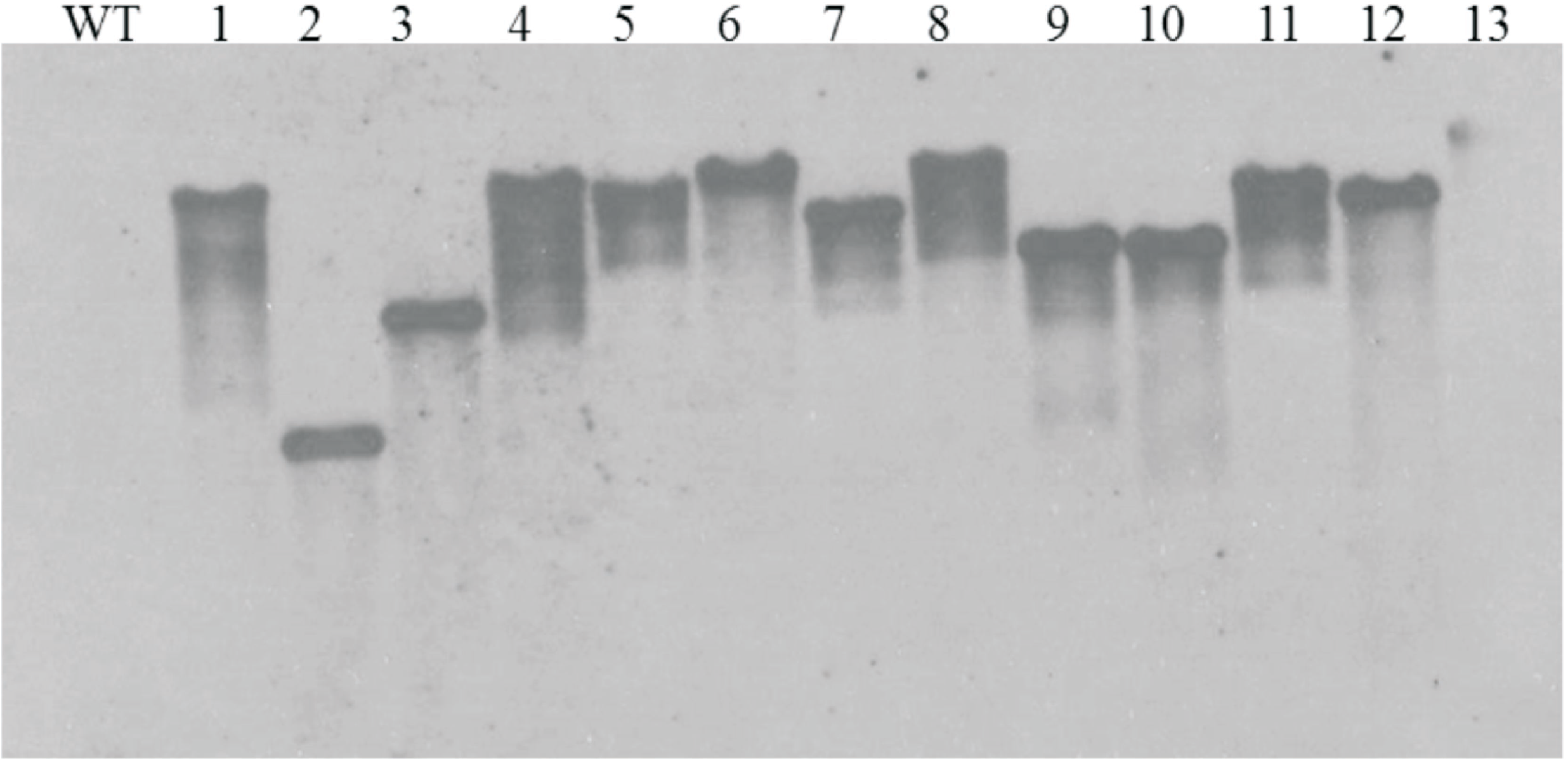
Southern blot analysis of PGPR2 INSeq library mutants. Genomic DNA of wild type PGPR2 (WT) and 13 INSeq mutants were digested with HindIII restriction enzyme and resolved on an 0.8% agarose gel. A gentamicin resistance gene was used as a probe revealed a single insertion and random integration of the transposon element.

**Fig. S3.**
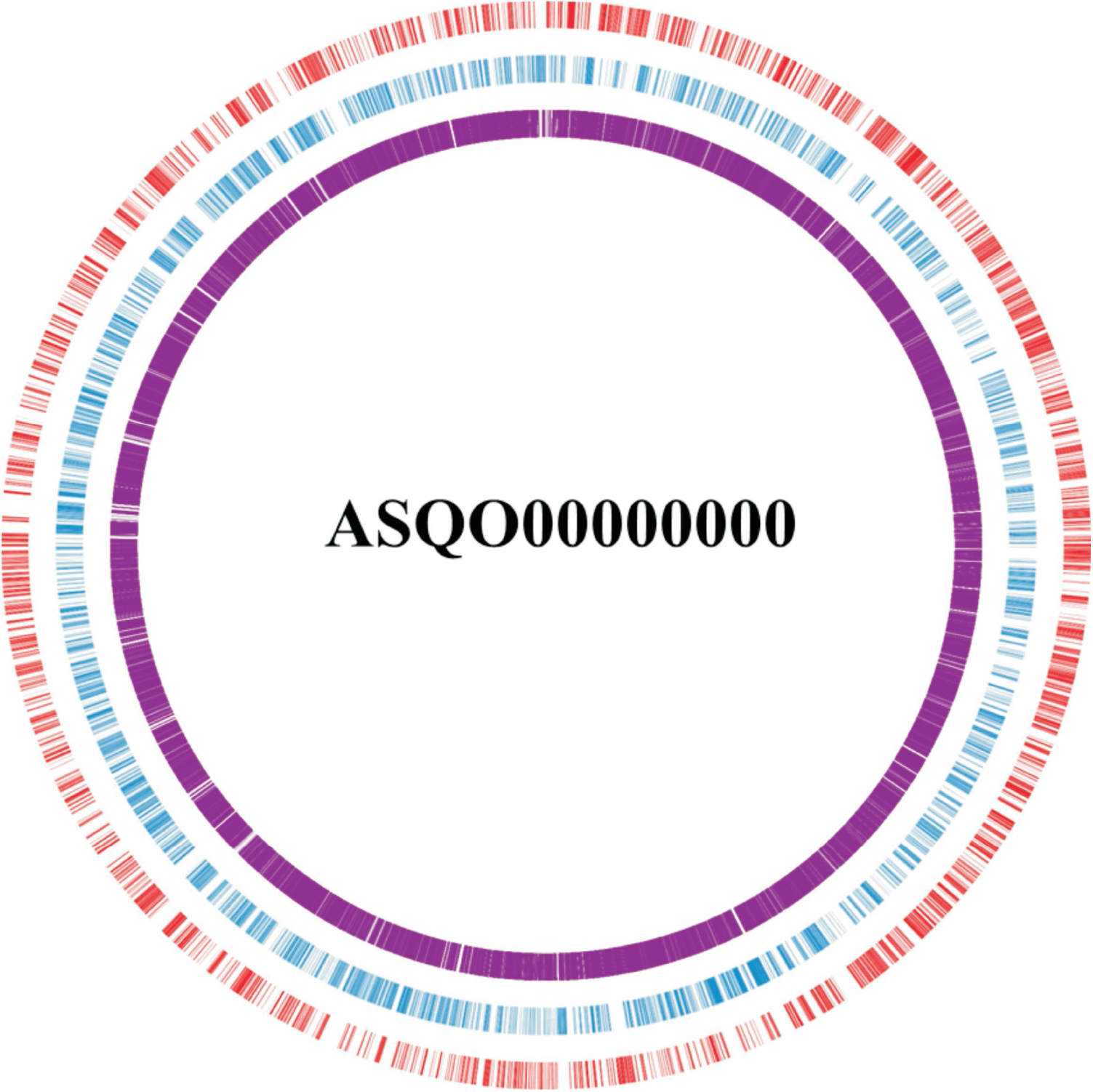

Genome map of *P. aeruginosa* PGPR2 showing transposon insertion sites. The outer circle represents the forward strand (red), the inner circle represents the reverse strand (blue), and the purple bars represent the transposon insertion sites.

**Fig. S4.**
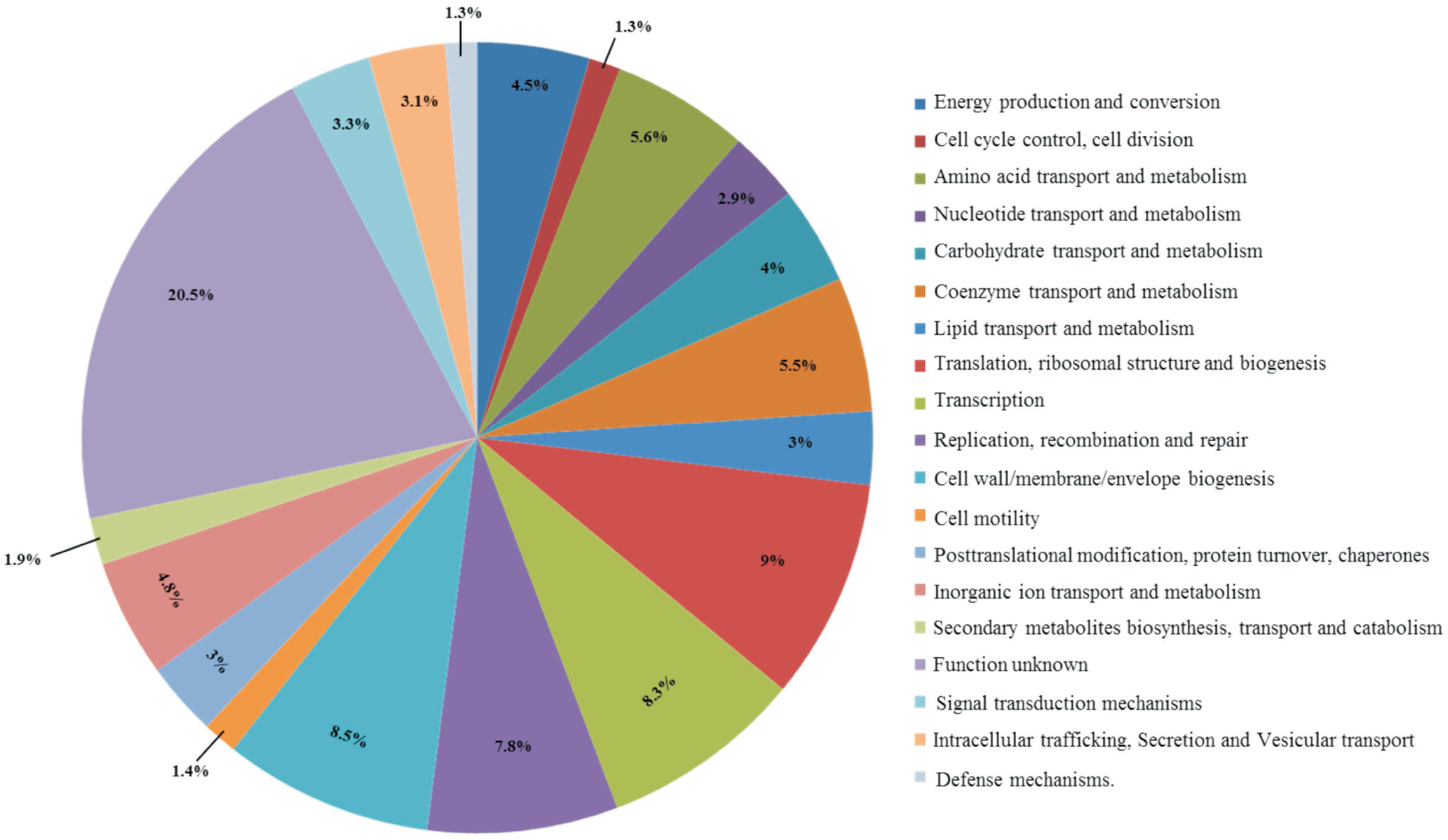

Functional categorization of essential genes responsible for growth of PGPR2. The functional classification was done according to protein annotation by COG database using WebMGA.

**Fig. S5.**
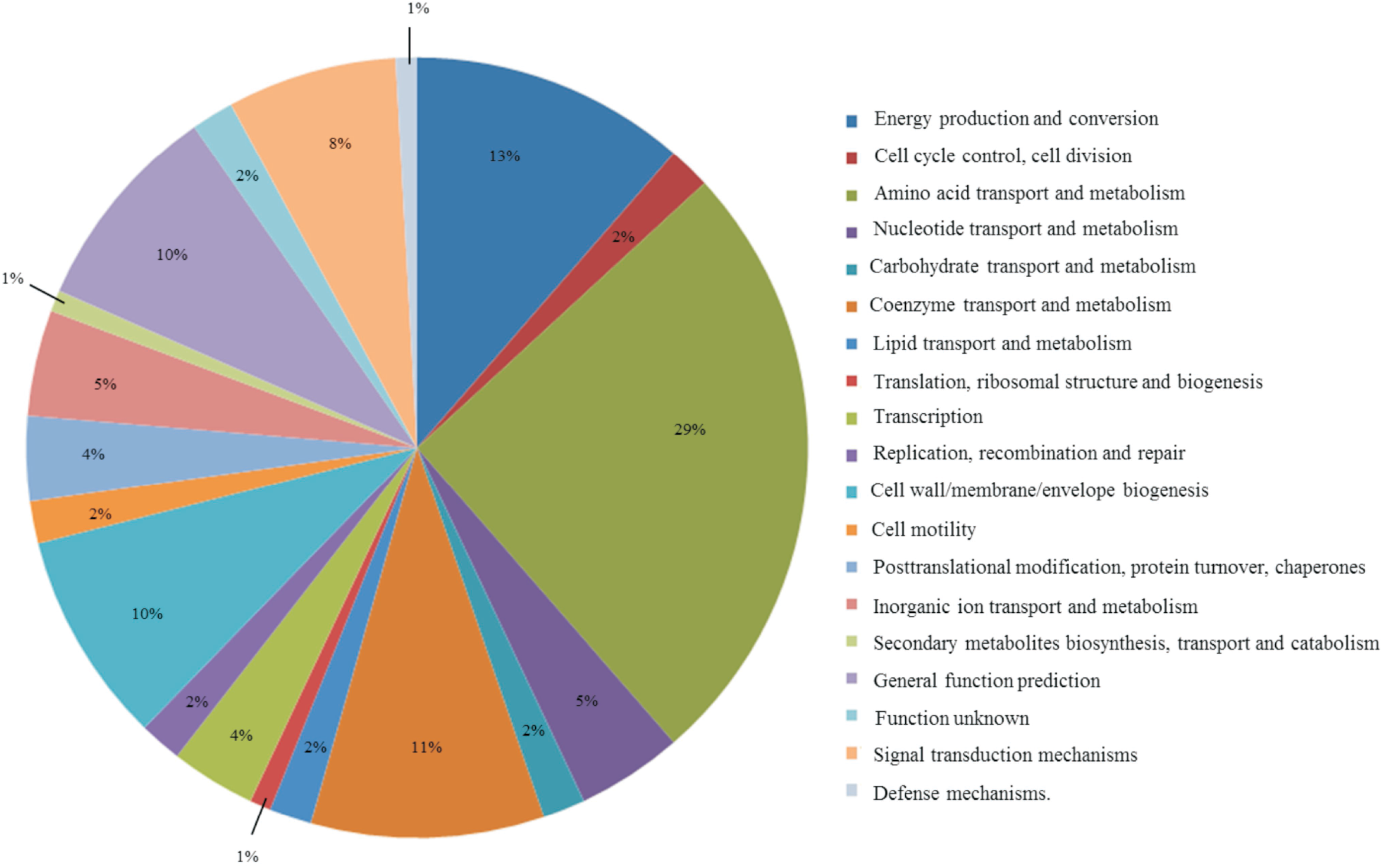

Functional categorization of genes responsible for the fitness of PGPR2 during corn root colonization. The functional classification was done according to protein annotation by COG database using WebMGA.

